## supplemental figures and tables for "Behavioral entrainment to rhythmic auditory stimulation can be modulated by tACS depending on the electrical stimulation field properties"

**This PDF file includes:**

Supplementary Text
Figs. S1 to S4
Tables S1 to S5

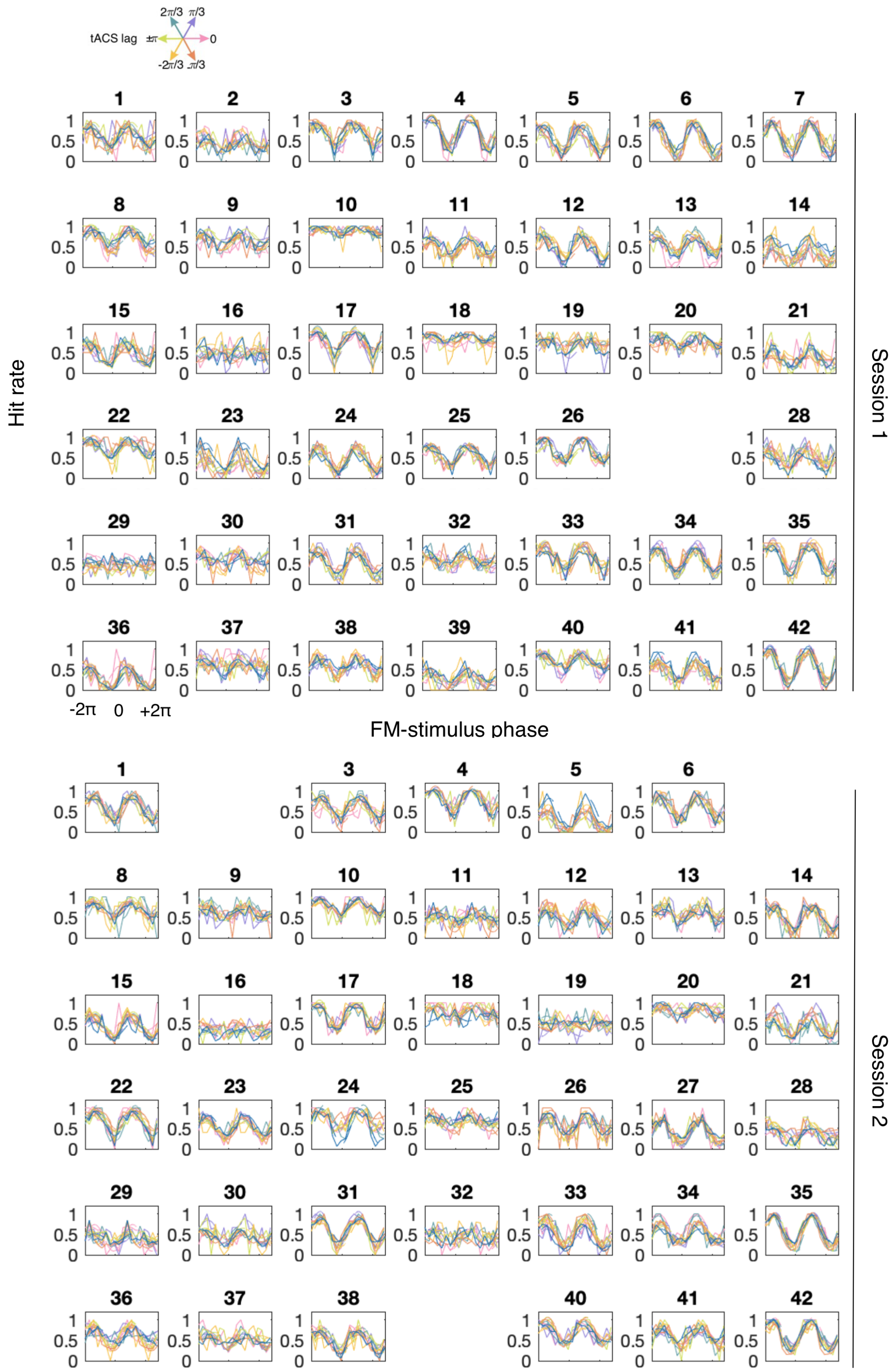


**Fig. S1. Gap detection performance.** Individual data for Fig. 3b. Hit rates as a function of FM-stimulus phase separated by tACS condition for both sessions. Colors follow the same coding as in the main Fig. 3b. Each plot represents a subject and session. Missing plots represent missing sessions.

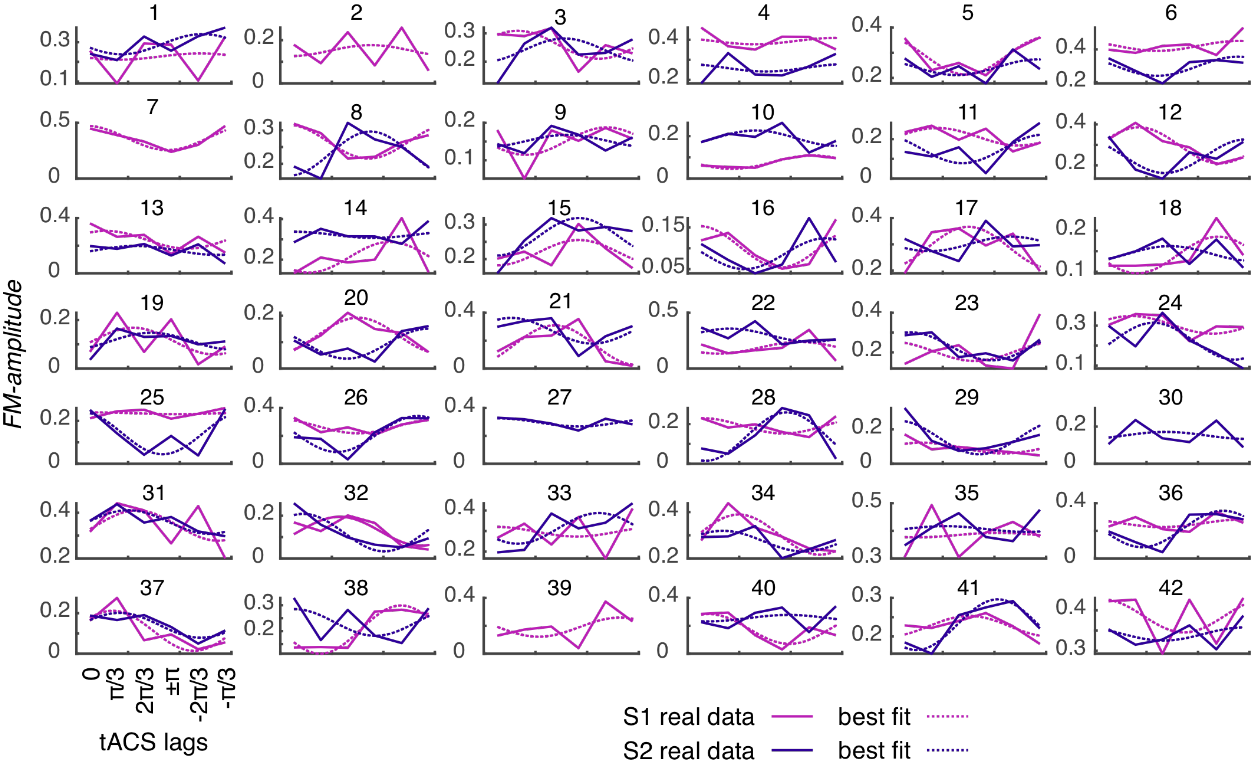


**Fig. S2. FM-amplitude as a function of tACS phase lag.** Individual data for Fig. 3e. Each plot shows data from a single participant. Solid lines show true data and dashed lines the cosine fit.

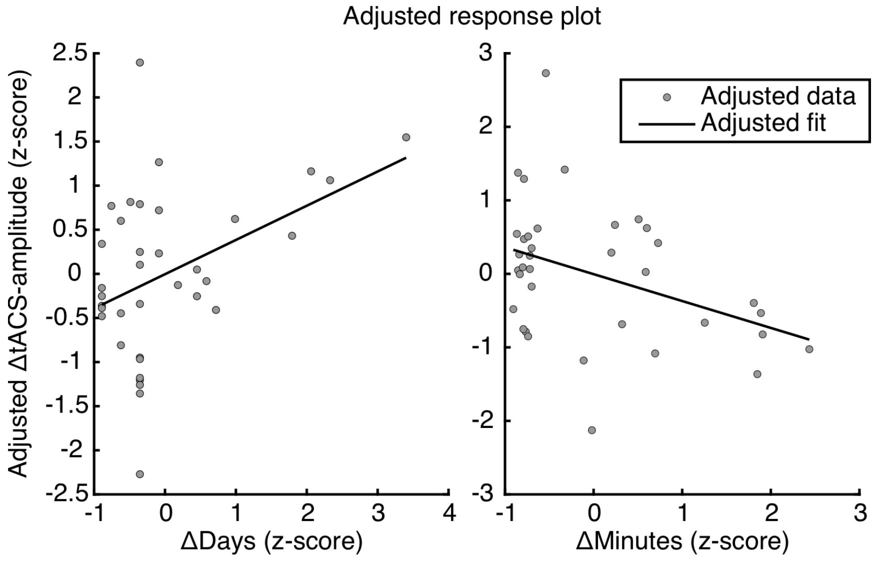


**Fig. S3. Predicting inter-session difference in tACS-amplitude.** Scatter plots show the change in tACS-amplitude between sessions (ΔtACS-amplitude. S1-S2) as a function of the number of days passed between sessions (ΔDays) and the time of the day absolute difference (ΔMinutes). Each dot represents a single participant. Black solid lines represent the adjusted fit.

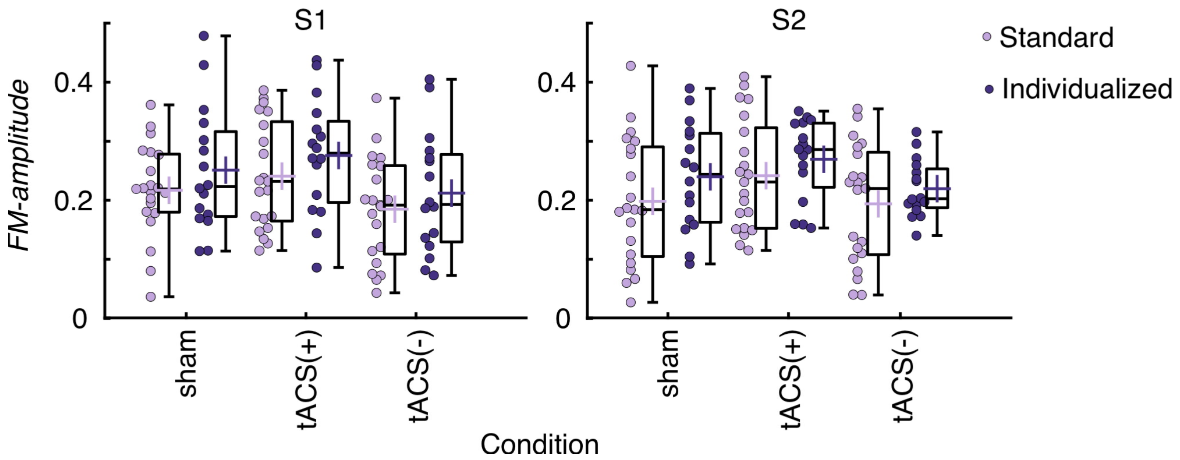

**Fig. S4. tACS effects separated by tACS montage.** For each session. FM-amplitude values estimated for sham. tACS(+) and tACS(-) are presented separated by tACS montage**.** Each dot represents a single participant. Box plots show median (vertical lines). mean (cross in the middle of the box). 25th and 75th percentiles (box edges) and extreme datapoints not considered outliers (+/– 2.7σ and 99.3 percentiles. whiskers). S1: session 1. S2: session 2.

**Table S1.** MNI center coordinates for target functional regions of interest for each subject

| Subject | Right hemisphere (x. y. z) | Left Hemisphere (x. y. z) |
|---|---|---|
| 1 | 56.328. -23.081. 5.419 | -53.954. -28.559. 8 |
| 2 | 50.722. -18.320. 7.010 | -54.714. -15.714. 6.286 |
| 3 | 61.478. -21.304. 7.565 | -56.820. -21.533. 5.072 |
| 4 | 57.482. -20.392. 1.396 | -53.491. -25.509. 4.541 |
| 5 | 58.336. -15.748. 4.899 | -54.254. -22.254. 5.559 |
| 6 | 56.226. -9.2260. 1.516 | -51.692. -10.462. 0.385 |
| 7 | 55.907. -25.389. 7.130 | -51.268. -29.895. 9.579 |
| 8 | 62.769. -22. 10.077 | -63.571. -23.571. 6.429 |
| 9 | 55.442. -17.731. 6.673 | -56.679. -22.643. 8.911 |
| 10 | 59.404. -29.527. 4.567 | -54.790. -25.559. 7.378 |
| 11 | 55.935. -15.613. 0.1610 | -51.500. -20.357. 1.286 |
| 12 | 54.150. -16.750. 3.500 | -57.343. -24.057. 9.114 |
| 13 | 52.759. -12.328. 2.724 | -48.783. -24.832. 9.081 |
| 14 | 58.917. -16.887. 3.702 | -54.884. -28.231. 6.787 |
| 15 | 52.340. -17.814. 3.633 | -50.180. -30.329. 10.273 |
| 16 | 54.368. -22.263. 3.737 | -48.278. -24.845. 7.320 |
| 17 | 58.643. -25.881. 9.238 | -54.026. -28.416. 7.675 |
| 18 | 57.471. -22.422. 7.922 | -54.475. -27.050. 8.307 |
| 19 | 58.098. -22.108. 5.647 | -53.211. -23.376. 7.193 |
| 20 | 62.803. -10.039. 3.618 | -49.769. -22.846. 9.462 |
| 21 | 60.239. -25.271. 3.401 | -56.011. -22.379. 4.629 |
| 22 | 52.548. -26.839. 13.226 | -57.951. -19.951. 5.659 |
| 23 | 50.070. -18.366. 5.127 | -50.029. -25.441. 8.971 |
| 24 | 55.563. -19.403. 7.672 | -54.150. -27.175. 6.950 |
| 25 | 58.549. -11.310. 1.028 | -52.122. -20.204. 5.571 |
| 26 | 53.714. -24.333. 6.952 | -51.435. -27.355. 6.419 |
| 27 | 59.127. -23.131. 7.138 | -51.872. -24.702. 6.830 |
| 28 | 47.600. -28.600. 5 | -55.721. -25.397. 9.588 |
| 29 | 59.524. -20.452. 3.595 | -60.577. -20.731. 6.154 |
| 30 | 54.923. -13.231. -1.462 | -55.589. -27.881. 9.144 |
| 31 | 53.910. -18.594. 4.075 | -52.071. -33.614. 10.614 |
| 32 | 57.364. -26.150. 8.196 | -61.533. -25.800. 7.467 |
| 33 | 55.762. -22.992. 7.053 | -53.278. -26.711. 9.422 |
| 34 | 54.862. -21.552. 8.862 | -50.090. -24.426. 7.761 |
| 35 | 58.468. -25.152. 3.949 | -55.136. -23.412. 6.038 |
| 36 | 58.347. -23.042. 6.627 | -53.846. -25.949. 7.360 |
| 37 | 62.104. -20.007. 5.582 | -54.661. -18.835. 4.009 |
| 38 | 57.733. -23.168. 5.426 | -51.351. -28.511. 10.218 |
| 39 | 59.427. -22.708. 6.6850 | -54.-27.407. 9.037 |

**Table S2.** Statistics for the mixed effects logistic regression models predicting single trial gap detection performance.

| | Formula | AIC | Δ AIC | BIC | LogLikelihood | Deviance |
|---|---|---|---|---|---|---|
| 1 | ***Gap detection* ~ 1 + sinFM + cosFM + (1 + sinFM + cosFM \| participant)** | **46560** | **0** | **46637** | **-23271** | **46542** |
| 2 | *Gap detection* ~ 1 + sintACS + costACS + sinFM + cosFM + (1 + sintACS + costACS + sinFM + cosFM \| participant) | 46566 | 6.203 | 46738 | -23263 | 46526 |
| 3 | *Gap detection* ~ 1 + sintACS*sinFM + costACS*sinFM + sintACS*cosFM + costACS*cosFM + (1 + sintACS:sinFM + costACS:sinFM + sintACS:cosFM + costACS:cosFM \| participant) | 47252 | 691.635 | 47457 | -23602 | 47204 |

Models are organized from smallest to highest AIC. Δ AIC relative to winning model. The winning model is also highlighted in bold. BIC: Bayesian information criterion.

**Table S3.** Statistics for the general linear models predicting inter-session difference on *tACS-amplitude*

| | Formula | AICc | Δ AICc* | $R^2$ | F** | p |
|---|---|---|---|---|---|---|
| **1** | ***ΔtACS-amplitude* ~ 1 + ΔDays + ΔMinutes** | **97.182** | **0** | **0.257** | **5.72** | **0.007** |
| 2 | *ΔtACS-amplitude* ~ 1 + Age + ΔDays + ΔMinutes | 97.310 | 0.129 | 0.306 | 4.69 | 0.008 |
| 3 | *ΔtACS-amplitude* ~ 1 + Age + ΔDays + ΔMinutes + Δgap size threshold | 97.629 | 0.447 | 0.35 | 4.18 | 0.008 |
| 4 | *ΔtACS-amplitude* ~ 1 + Age + ΔDays + ΔMinutes + Δgap size threshold + Montage | 100.047 | 2.866 | 0.359 | 3.36 | 0.016 |
| 5 | *ΔtACS-amplitude* ~ 1 ΔDays | 100.289 | 3.107 | 0.135 | 5.31 | 0.027 |
| 6 | *ΔtACS-amplitude* ~ 1 + ΔMinutes | 101.269 | 4.087 | 0.111 | 4.25 | 0.047 |
| 7 | *ΔtACS-amplitude* ~ 1 + Gender + Age + ΔDays + ΔMinutes + Δgap size threshold + Montage | 102.803 | 5.621 | 0.365 | 3.78 | 0.029 |

Models are organized from smallest to highest AICc. * Δ AICc relative to winning model. ** F model vs. constant. The winning model is also highlighted in bold.

**Table S4.** Statistics for the general linear models predicting inter-session absolute circular distance on *tACS-phase*

| | Formula | AICc | Δ AICc* | $R^2$ | F** | p |
|---|---|---|---|---|---|---|
| 1 | \|*circ_distance*\| ~ 1 + gender | 102.416 | 0 | 0.082 | 3.05 | 0.09 |
| 2 | \|*circ_distance*\| ~ 1 + gender + Δgap size threshold | 102.703 | 0.288 | 0.134 | 2.56 | 0.092 |
| 3 | \|*circ_distance*\| ~ 1 + gender + ΔMinutes + Δgap size threshold | 103.285 | 0.869 | 0.18 | 2.35 | 0.091 |
| 4 | \|*circ_distance*\| ~ 1 + gender + Age + ΔMinutes + Δgap size threshold | 104.498 | 2.083 | 0.214 | 2.11 | 0.104 |
| 5 | \|*circ_distance*\| ~ 1 + gender + Age + ΔMinutes + Δgap size threshold + Montage | 106.564 | 4.148 | 0.232 | 1.81 | 0.141 |
| 6 | \|*circ_distance*\| ~ 1 + gender + Age + ΔDays + ΔMinutes + Δgap size threshold + Montage | 109.663 | 7.247 | 0.232 | 1.46 | 0.228 |

Models are organized from smallest to highest AICc. * Δ AICc relative to winning model. ** F model vs. constant. No model was significant relative to a constant model.

**Table S5.** Statistics for the general linear models predicting tACS effects

| | Formula | AICc | Δ AICc* | $R^2$ | F** | p |
|---|---|---|---|---|---|---|
| 1 | ***tACS_amplitude* ~1 + Dist2Peak*Normal_E_field + Normal_E_field*Focality** | **94.429** | **0** | **0.378** | **3.406** | **0.016** |
| 2 | tACS_amplitude ~1 + Corr2BOLD*Normal_E_field_ROI + Normal_E_field_ROI*Focality | 95.691 | 1.262 | 0.355 | 3.078 | 0.024 |
| 3 | tACS_amplitude ~1 + Corr2BOLD*E_field_ROI + E_field_ROI*Focality | 95.853 | 1.423 | 0.352 | 3.036 | 0.026 |
| 4 | tACS_amplitude ~1 + Corr2BOLD*Normal_E_field + Normal_E_field*Focality | 96.719 | 2.289 | 0.335 | 2.819 | 0.035 |
| 5 | tACS_amplitude ~1 + Dist2Peak*E_field + E_field*Focality | 97.156 | 2.727 | 0.326 | 2.712 | 0.040 |
| 6 | tACS_amplitude ~1 + Dist2Peak*Normal_E_field + Dist2Peak*Focality + Normal_E_field*Focality | 97.535 | 3.105 | 0.380 | 2.757 | 0.032 |
| 7 | tACS_amplitude ~1 + Corr2BOLD*E_field + E_field*Focality | 97.727 | 3.297 | 0.315 | 2.574 | 0.049 |
| 8 | tACS_amplitude ~1 + Focality + Corr2BOLD*Normal_E_field_ROI | 97.738 | 3.308 | 0.252 | 2.444 | 0.069 |
| 9 | tACS_amplitude ~1 + Dist2Peak + Normal_E_field_ROI*Focality | 97.901 | 3.472 | 0.249 | 2.398 | 0.073 |
| 10 | tACS_amplitude ~1 + Dist2Peak*Normal_E_field_ROI + Normal_E_field_ROI*Focality | 98.022 | 3.592 | 0.309 | 2.503 | 0.054 |
| 11 | tACS_amplitude ~1 + Corr2BOLD + Normal_E_field_ROI*Focality | 98.267 | 3.838 | 0.240 | 2.294 | 0.083 |
| 12 | tACS_amplitude ~1 + Dist2Peak*Normal_E_field_ROI + Dist2Peak*Focality + Normal_E_field_ROI*Focality | 98.347 | 3.917 | 0.365 | 2.585 | 0.041 |
| 13 | tACS_amplitude ~1 + Corr2BOLD*Normal_E_field_ROI + Corr2BOLD*Focality + Normal_E_field_ROI*Focality | 98.629 | 4.200 | 0.359 | 2.527 | 0.045 |
| 14 | tACS_amplitude ~1 + Focality + Corr2BOLD*E_field | 98.689 | 4.260 | 0.231 | 2.177 | 0.097 |
| 15 | tACS_amplitude ~1 + Corr2BOLD*E_field_ROI + Corr2BOLD*Focality + E_field_ROI*Focality | 98.858 | 4.429 | 0.355 | 2.480 | 0.048 |
| 16 | tACS_amplitude ~1 + Corr2BOLD*Normal_E_field + Corr2BOLD*Focality + Normal_E_field*Focality | 99.118 | 4.689 | 0.350 | 2.426 | 0.053 |
| 17 | tACS_amplitude ~1 + Focality + Corr2BOLD*Normal_E_field | 99.319 | 4.889 | 0.217 | 2.004 | 0.120 |
| 18 | tACS_amplitude ~1 + Dist2Peak*E_field + Dist2Peak*Focality + E_field*Focality | 99.423 | 4.993 | 0.344 | 2.365 | 0.058 |
| 19 | tACS_amplitude ~1 + Dist2Peak + Normal_E_field*Focality | 99.877 | 5.448 | 0.204 | 1.853 | 0.146 |
| 20 | tACS_amplitude ~1 + Focality + Corr2BOLD*E_field_ROI | 99.889 | 5.459 | 0.203 | 1.850 | 0.146 |
| 21 | tACS_amplitude ~1 + Corr2BOLD + Normal_E_field*Focality | 99.889 | 5.460 | 0.203 | 1.850 | 0.146 |
| 22 | tACS_amplitude ~1 + Dist2Peak + Normal_E_field_ROI + Focality | 99.912 | 5.483 | 0.135 | 1.564 | 0.219 |
| 23 | tACS_amplitude ~1 + Corr2BOLD + Normal_E_field_ROI + Focality | 100.18 | 5.753 | 0.128 | 1.472 | 0.242 |
| 24 | tACS_amplitude ~1 + Sham + Corr2BOLD*E_field_ROI + Corr2BOLD*Focality + E_field_ROI*Focality | 100.289 | 5.860 | 0.392 | 2.399 | 0.049 |
| 25 | tACS_amplitude ~1 + Corr2BOLD*E_field + Corr2BOLD*Focality + E_field*Focality | 100.38 | 5.948 | 0.326 | 2.174 | 0.077 |
| 26 | tACS_amplitude ~1 + Dist2Peak + E_field*Focality | 100.38 | 5.954 | 0.192 | 1.719 | 0.173 |
| 27 | tACS_amplitude ~1 + Corr2BOLD + E_field*Focality | 100.42 | 5.988 | 0.191 | 1.710 | 0.175 |

| | | | | | | |
|---|---|---|---|---|---|---|
| 28 | tACS_amplitude ~1 + BOLDbetaMean + Dist2Peak*Normal_E_field + Dist2Peak*Focality + Normal_E_field*Focality | 100.572 | 6.143 | 0.387 | 2.349 | 0.053 |
| 29 | tACS_amplitude ~1 + Dist2Peak + E_field_ROI*Focality | 100.66 | 6.232 | 0.185 | 1.646 | 0.190 |
| 30 | tACS_amplitude ~1 + Corr2BOLD + E_field_ROI*Focality | 100.70 | 6.272 | 0.184 | 1.635 | 0.192 |
| 31 | 'tACS_amplitude ~1 + Sham + Corr2BOLD*Normal_E_field_ROI + Corr2BOLD*Focality + Normal_E_field_ROI*Focality' | 100.766 | 6.337 | 0.384 | 2.314 | 0.056 |
| 32 | tACS_amplitude ~1 + Sham + Dist2Peak*Normal_E_field_ROI + Dist2Peak*Focality + Normal_E_field_ROI*Focality | 100.845 | 6.415 | 0.382 | 2.300 | 0.058 |
| 33 | tACS_amplitude ~1 + Sham + Dist2Peak*Normal_E_field + Dist2Peak*Focality + Normal_E_field*Focality | 100.969 | 6.540 | 0.380 | 2.278 | 0.060 |
| 34 | tACS_amplitude ~1 + Dist2Peak*E_field_ROI + E_field_ROI*Focality | 100.99 | 6.564 | 0.246 | 1.825 | 0.140 |
| 35 | tACS_amplitude ~1 + Dist2Peak + Normal_E_field + Focality | 101.41 | 6.984 | 0.096 | 1.064 | 0.379 |
| 36 | tACS_amplitude ~1 + Normal_E_field_ROI + Dist2Peak*Focality | 101.42 | 6.986 | 0.167 | 1.450 | 0.243 |
| 37 | tACS_amplitude ~1 + BOLDbetaMean + Dist2Peak*Normal_E_field_ROI + Dist2Peak*Focality + Normal_E_field_ROI*Focality | 101.547 | 7.118 | 0.370 | 2.177 | 0.070 |
| 38 | tACS_amplitude ~1 + Dist2Peak*E_field_ROI + Dist2Peak*Focality + E_field_ROI*Focality | 101.63 | 7.200 | 0.300 | 1.933 | 0.112 |
| 39 | tACS_amplitude ~1 + Dist2Peak + E_field_ROI + Focality | 101.65 | 7.220 | 0.090 | 0.988 | 0.412 |
| 40 | tACS_amplitude ~1 + Normal_E_field_ROI + Corr2BOLD*Focality | 101.67 | 7.242 | 0.160 | 1.385 | 0.264 |
| 41 | tACS_amplitude ~1 + Dist2Peak + E_field + Focality | 101.70 | 7.269 | 0.089 | 0.972 | 0.419 |
| 42 | tACS_amplitude ~1 + Corr2BOLD + Normal_E_field + Focality | 101.97 | 7.538 | 0.081 | 0.885 | 0.460 |
| 43 | tACS_amplitude ~1 + Sham + Dist2Peak*E_field + Dist2Peak*Focality + E_field*Focality | 102.034 | 7.604 | 0.360 | 2.094 | 0.081 |
| 44 | tACS_amplitude ~1 + BOLDbetaMean + Corr2BOLD*E_field_ROI + Corr2BOLD*Focality + E_field_ROI*Focality | 102.06124410104 | 7.63183993644338 | 0.3599588489437778 | 2.0889125771932 | 0.0812024639062292 |
| 45 | tACS_amplitude ~1 + BOLDbetaMean + Corr2BOLD*Normal_E_field_ROI + Corr2BOLD*Focality + Normal_E_field_ROI*Focality | 102.065 | 7.636 | 0.360 | 2.088 | 0.081 |
| 46 | tACS_amplitude ~1 + Corr2BOLD + E_field_ROI + Focality | 102.07 | 7.642 | 0.079 | 0.852 | 0.476 |
| 47 | tACS_amplitude ~1 + Sham + Corr2BOLD*Normal_E_field + Corr2BOLD*Focality + Normal_E_field*Focality | 102.071 | 7.642 | 0.360 | 2.087 | 0.081 |
| 48 | tACS_amplitude ~1 + Corr2BOLD + E_field + Focality | 102.20 | 7.767 | 0.075 | 0.812 | 0.497 |
| 49 | tACS_amplitude ~1 + Focality + Dist2Peak*Normal_E_field_ROI | 102.21 | 7.780 | 0.147 | 1.250 | 0.312 |
| 50 | tACS_amplitude ~1 + BOLDbetaMean + Dist2Peak*E_field + Dist2Peak*Focality + E_field*Focality | 102.274 | 7.845 | 0.356 | 2.053 | 0.086 |
| 51 | tACS_amplitude ~1 + Focality + Dist2Peak*Normal_E_field | 102.40 | 7.974 | 0.142 | 1.201 | 0.331 |

| | | | | | | |
|---|---|---|---|---|---|---|
| 52 | tACS_amplitude ~1 + BOLDbetaMean + Corr2BOLD*Normal_E_field + Corr2BOLD*Focality + Normal_E_field*Focality | 102.424 | 7.994 | 0.353 | 2.027 | 0.090 |
| 53 | tACS_amplitude ~1 + Sham + Corr2BOLD*E_field + Corr2BOLD*Focality + E_field*Focality | 102.481 | 8.052 | 0.352 | 2.018 | 0.091 |
| 54 | tACS_amplitude ~1 + Focality + Dist2Peak*E_field | 102.50 | 8.073 | 0.140 | 1.177 | 0.342 |
| 55 | tACS_amplitude ~1 + Dist2Peak*E_field + Dist2Peak*Focality + E_field*Focality + Dist2Peak:E_field:Focality | 102.57 | 8.143 | 0.350 | 2.002 | 0.094 |
| 56 | tACS_amplitude ~1 + Normal_E_field + Dist2Peak*Focality | 102.58 | 8.150 | 0.138 | 1.158 | 0.350 |
| 57 | tACS_amplitude ~1 + E_field_ROI + Dist2Peak*Focality | 102.78 | 8.353 | 0.133 | 1.107 | 0.372 |
| 58 | tACS_amplitude ~1 + E_field + Dist2Peak*Focality | 102.90 | 8.466 | 0.130 | 1.080 | 0.385825 |
| 59 | tACS_amplitude ~1 + Sham + Dist2Peak*E_field_ROI + Dist2Peak*Focality + E_field_ROI*Focality | 103.589 | 9.160 | 0.331 | 1.834 | 0.123 |
| 60 | tACS_amplitude ~1 + BOLDbetaMean + Corr2BOLD*E_field + Corr2BOLD*Focality + E_field*Focality | 103.761 | 9.332 | 0.327 | 1.806 | 0.129 |
| 61 | tACS_amplitude ~1 + E_field_ROI + Corr2BOLD*Focality | 103.84 | 9.408 | 0.105 | 0.852 | 0.504 |
| 62 | tACS_amplitude ~1 + Normal_E_field + Corr2BOLD*Focality | 103.90 | 9.473 | 0.103 | 0.837 | 0.513 |
| 63 | tACS_amplitude ~1 + Sham + BOLDbetaMean + Corr2BOLD*E_field_ROI + Corr2BOLD*Focality + E_field_ROI*Focality | 103.975 | 9.546 | 0.393 | 2.027 | 0.085 |
| 64 | tACS_amplitude ~1 + Focality + Dist2Peak*E_field_ROI | 104.11 | 9.676 | 0.098 | 0.789 | 0.542 |
| 65 | tACS_amplitude ~1 + E_field + Corr2BOLD*Focality | 104.17 | 9.737 | 0.096 | 0.774 | 0.551 |
| 66 | tACS_amplitude ~1 + Sham + BOLDbetaMean + Dist2Peak*Normal_E_field + Dist2Peak*Focality + Normal_E_field*Focality | 104.31 | 9.877 | 0.387 | 1.977 | 0.092 |
| 67 | tACS_amplitude ~1 + Sham + BOLDbetaMean + Dist2Peak*Normal_E_field_ROI + Dist2Peak*Focality + Normal_E_field_ROI*Focality | 104.439 | 10.010 | 0.385 | 1.957 | 0.095 |
| 68 | tACS_amplitude ~1 + Sham + BOLDbetaMean + Corr2BOLD*Normal_E_field_ROI + Corr2BOLD*Focality + Normal_E_field_ROI*Focality | 104.506 | 10.076 | 0.384 | 1.947 | 0.097 |
| 69 | tACS_amplitude ~1 + BOLDbetaMean + Dist2Peak*E_field_ROI + Dist2Peak*Focality + E_field_ROI*Focality | 104.526 | 10.096 | 0.312 | 1.683 | 0.157 |
| 70 | tACS_amplitude ~1 + Dist2Peak + Normal_E_field_ROI + Focality + Sham + BOLDbetaMean | 104.87 | 10.44 | 0.155 | 1.025 | 0.422 |
| 71 | tACS_amplitude ~1 + Corr2BOLD + Normal_E_field_ROI + Focality + Sham + BOLDbetaMean | 105.07 | 10.64 | 0.150 | 0.986 | 0.444 |
| 72 | tACS_amplitude ~1 + Sham + BOLDbetaMean + Dist2Peak*E_field + Dist2Peak*Focality + E_field*Focality | 105.285 | 10.856 | 0.370 | 1.832 | 0.118 |
| 73 | tACS_amplitude ~1 + Sham + BOLDbetaMean + Corr2BOLD*Normal_E_field + Corr2BOLD*Focality + Normal_E_field*Focality | 105.694 | 11.265 | 0.362 | 1.773 | 0.131 |
| 74 | tACS_amplitude ~1 + Sham + BOLDbetaMean + Corr2BOLD*E_field + Corr2BOLD*Focality + E_field*Focality | 106.201 | 11.772 | 0.352 | 1.700 | 0.148 |

| | | | | | | |
|---|---|---|---|---|---|---|
| 75 | tACS_amplitude ~1 + Dist2Peak + Normal_E_field + Focality + Sham + BOLDbetaMean | 106.215 | 11.785 | 0.121 | 0.768 | 0.581 |
| 76 | tACS_amplitude ~1 + Dist2Peak + E_field + Focality + Sham + BOLDbetaMean | 106.693 | 12.264 | 0.108 | 0.679 | 0.643 |
| 77 | tACS_amplitude ~1 + Corr2BOLD + Normal_E_field + Focality + Sham + BOLDbetaMean | 106.693 | 12.264 | 0.108 | 0.679 | 0.643 |
| 78 | tACS_amplitude ~1 + Dist2Peak + E_field_ROI + Focality + Sham + BOLDbetaMean' | 106.752 | 12.322 | 0.107 | 0.668 | 0.651 |
| 79 | tACS_amplitude ~1 + Sham + BOLDbetaMean + Dist2Peak*E_field_ROI + Dist2Peak*Focality + E_field_ROI*Focality | 106.965 | 12.535 | 0.338 | 1.593 | 0.177 |
| 80 | tACS_amplitude ~1 + Corr2BOLD + E_field_ROI + Focality + Sham + BOLDbetaMean | 107.112 | 12.681 | 0.097 | 0.602 | 0.699 |
| 81 | tACS_amplitude ~1 + Corr2BOLD + E_field + Focality + Sham + BOLDbetaMean | 107.112 | 12.683 | 0.097 | 0.602 | 0.699 |

Models are organized from smallest to highest AICc. * Δ AICc relative to winning model. ** F model vs. constant. The winning model is also highlighted in bold.
